# Spatially heterogeneous responses of planktonic foraminifera assemblages over 700,000 years of climate change

**DOI:** 10.1101/2024.03.08.584139

**Authors:** Gregor H. Mathes, Carl J. Reddin, Wolfgang Kiessling, Gawain S. Antell, Erin E. Saupe, Manuel J. Steinbauer

## Abstract

**Aim:** To determine the degree to which assemblages of planktonic foraminifera track thermal conditions.

**Location:** The world’s oceans.

**Time period:** The last 700,000 years of glacial-interglacial cycles.

**Major taxa studied:** Planktonic foraminifera.

**Methods:** We investigate assemblage dynamics in planktonic foraminifera in response to temperature changes using a global dataset of Quaternary planktonic foraminifera, together with a coupled Atmosphere–Ocean General Circulation Model (AOGCM) at 8,000-year resolution. We use ‘thermal deviance’ to assess assemblage responses to climate change, defined as the difference between the temperature at a given location and the bio-indicated temperature (i.e., the abundance-weighted average of estimated temperature optima for the species present).

**Results:** Assemblages generally tracked annual mean temperature changes through compositional turnover, but large thermal deviances are evident under certain conditions. The coldest-adapted species persisted in polar regions during warming but were not joined by additional immigrants, resulting in decreased assemblage turnover with warming. The warmest-adapted species persisted in equatorial regions during cooling. Assemblages at mid latitudes closely tracked temperature cooling and showed a modest increase in thermal deviance with warming.

**Main conclusions:** Planktonic foraminiferal assemblages were generally able to track or endure temperature changes: as climate warmed or cooled, bio-indicated temperature also became warmer or cooler, although to a variable degree. At polar sites under warming and at equatorial sites under cooling, the change in temperature predicted from assemblage composition was less than, or even opposite to, expectations based on estimated environmental change. Nevertheless, all species survived the accumulation of thermal deviance—a result that highlights the resilience and inertia of planktonic foraminifera on an assemblage level to the last 700,000 years of climate change, which might be facilitated by broad thermal tolerances or depth shifts.

## Introduction

Anthropogenic climate change is threatening marine ecosystems (Cooley et al., 2022). A common response of marine species to changing climate is shifts in distribution to track suitable conditions (Chen, Hill, Ohlemüller, Roy, & Thomas, 2011; Lenoir et al., 2020; Pinsky, Worm, Fogarty, Sarmiento, & Levin, 2013; Poloczanska et al., 2013). However, the degree to which marine species keep pace with current rates of climate change via dispersal remains uncertain (García Molinos et al., 2016; Munday, Warner, Monro, Pandolfi, & Marshall, 2013). Although individual species of marine ectotherms are projected to closely track their thermal limits (Sunday, Bates, & Dulvy, 2012), assemblages are unlikely to respond cohesively to climate change (García Molinos et al., 2016; Graham et al., 1996; Walther et al., 2002), which can be attributed to the differential needs and tolerances of individual species across multiple abiotic parameters (Strack, Jonkers, C Rillo, Hillebrand, & Kucera, 2022). The individualistic responses of species to climate change may make certain regions or populations within assemblages more vulnerable to extirpation and global extinction (Reddin, Aberhan, Raja, & Kocsis, 2022; Stuart-Smith, Edgar, Barrett, Kininmonth, & Bates, 2015). This vulnerability can be indicated by a discrepancy between ambient temperatures and the conditions preferred by individual species within assemblages (i.e., thermal deviance; Devictor et al., 2012; Menéndez et al., 2006; Svenning & Sandel, 2013). However, the paucity of long-term studies prevent a full understanding of compositional changes in response to climate change (Rosenzweig et al., 2008). Mismatches between existing conditions and average temperature preferences of the assemblage constituents are studied more in terrestrial than in marine ecosystems (Sunday, Bates, & Dulvy, 2011). This differential focus may be warranted due to the greater magnitude and incidence of dispersal barriers on land and the lower vagility of many terrestrial species, which cannot disperse on wind or water currents (Burgess, Baskett, Grosberg, Morgan, & Strathmann, 2016). However, conclusions from global change biology studies in the terrestrial realm may not always translate to the marine realm (Knapp et al., 2017).

The exceptional fossil record of planktonic foraminifera renders them an ideal system to investigate thermal deviances in the ocean. Planktonic foraminifera are a near-ubiquitous component of the marine zooplankton. Their calcite shells (tests) are well-preserved in seafloor sediments, allowing for the composition of marine assemblages to be quantified at high temporal and spatial resolution. The tests of planktonic foraminifera have been used to investigate climatic and ecological changes in both modern and fossil systems (Antell, Fenton, Valdes, & Saupe, 2021; Ezard, Aze, Pearson, & Purvis, 2011; Jonkers, Hillebrand, & Kucera, 2019; Morard et al., 2015; Yasuhara, Okahashi, Cronin, Rasmussen, & Hunt, 2014; Yasuhara, Wei, et al., 2020).

Modern planktonic foraminifera assemblages have changed in community structure as a response to anthropogenic climate warming (Jonkers et al., 2019; Yasuhara, Huang, et al., 2020). The fossil record provides complementary data on how species have responded to various magnitudes, rates, and directions of climate change. Since unique combinations of climatic variables existed in the past (Williams & Jackson, 2007) and are projected to emerge in the future (Burke et al., 2018; Lotterhos, Láruson, & Jiang, 2021; Williams, Jackson, & Kutzbach, 2007), critical information can be obtained from the fossil record on biotic responses to climate conditions outside modern human experience. Most studies using the fossil record, however, analyse temporal resolutions of 10^6^ to 10^7^ years, above the timescales relevant to anthropogenic changes, or are restricted to a few discrete historical time steps. The continuous nature of the planktonic foraminifera fossil record provides an opportunity to study the impacts of climate change on spatio-temporal assemblage patterns at a temporal scale relevant for modern-day global change biology.

Here we investigate the degree to which planktonic foraminifera assemblages changed over the past 700,000 years of glacial-interglacial cycles. We base our analysis on a global dataset of Quaternary planktonic foraminifera records at 8-thousand-year (ka) resolution, coupled with surface temperature data from an Atmosphere–Ocean General Circulation Model (AOGCM). For each foraminiferal assemblage, we calculated thermal deviance as the difference between the modelled temperature at the location and the “bio-indicated temperature” for the local assemblage, quantified as the abundance-weighted average across estimated temperature optima of the individual species’ distribution (Fig. 1). We then used generalised additive and generalised linear models to quantify thermal deviance as a function of temperature change both globally and within latitudinal bands (low, mid, and high latitudes). We further link these patterns to changes in compositional turnover, latitudinal diversity, and the probability of local extinction (extirpation) in response to climate change.

**Figure 1:**
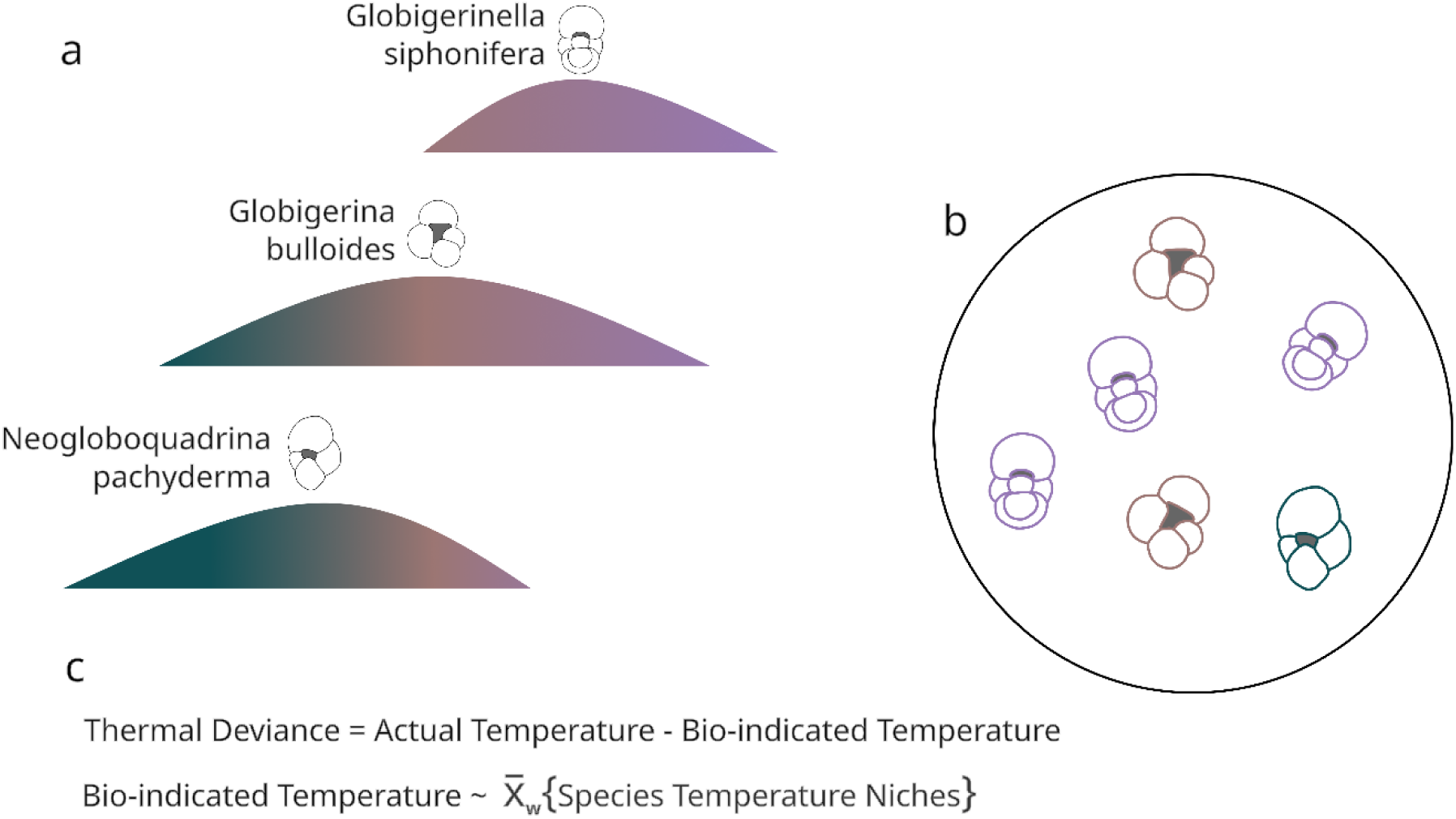
Calculation of the thermal deviance of a planktonic foraminifera assemblage. (a) Individual planktonic foraminifera species occupy a specific temperature niche. This temperature niche can be estimated for each species by documenting the response of fossil foraminifera species along the univariate axis of mean annual temperature. (b) An assemblage of planktonic foraminifera consists of various species, each displaying a characteristic temperature niche. The temperature of an assemblage (‘bio-indicated temperature’) can be estimated by weighted-averaging the temperature niches of individual species via an ecological transfer function (^x̅^ _w_), which takes both species composition and relative abundance of species into account. (c) The thermal deviance of the assemblage is then quantified as the difference between the modelled temperature at the location of the assemblage and the bio-indicated temperature of the assemblage.

We expected fossil planktonic foraminifera assemblages to closely track climate change given the ecologically broad temporal scale of our study, and the low number of barriers and high dispersal potential for marine plankton.

## Material and Methods

### Fossil data

Foraminiferal occurrence data over the last 700 ka are from Antell et al. (2021), derived from Fenton et al. (2021). Data from these sources consist of curated and taxonomically-harmonised planktonic foraminifera occurrences with recent age models. We removed occurrences from 0-100 m depth, following Antell et al. (2021), since these fossils have a higher probability of being transported from their life position. We retained those sites with information about the relative abundance of individual species and with occurrences in at least 10 consecutive time intervals globally. The final dataset contained 55 species with 38,441 occurrences (see Supporting Information Appendix S1 Table S1).

Occurrence records were binned into 88 time intervals of 8 ka resolution, from the recent subfossil record to 704 ka, following Antell et al. (2021). The chosen bin duration allowed for imprecision in fossil ages, typically on the order of a few thousand years (Martin, 1999), and agrees with the resolution of the coupled Atmosphere–Ocean General Circulation Model outputs (4 ka) and with general time-averaging expected in fossil assemblages of planktonic foraminifera assemblages (see Martin, 1999). The bin duration also ensures ecological dynamics were captured at a higher resolution than the fluctuations from glacial minimum to interglacial maximum.

We use the term ‘assemblage’ to refer to all species present within a sediment sample (i.e., a single location on Earth) in a given time bin of 8,000-year duration. The final dataset contains 2,607 assemblages (see Appendix S1 Table S2).

### Temperature data

Temperature is likely the single most important explanatory variable structuring the geographic distributions of planktonic foraminifera (Fenton, Pearson, Dunkley Jones, & Purvis, 2016; Yasuhara, Wei, et al., 2020). One way to reconstruct temperatures of prehistoric oceans is using coupled Atmosphere–Ocean General Circulation Models (AOGCM). We used mean annual surface temperature (MAT) from Antell et al. (2021), extracted from the UK Met Office Hadley Centre Coupled Model (Valdes et al., 2017). This model family is known to perform well in capturing mean climate state (Flato et al., 2014; Reichler & Kim, 2008; Valdes et al., 2017). The precise model configuration is documented in Valdez et al. (2017). Climate data were downscaled to 1.25° × 1.25° horizontal resolution at 20 vertical ocean levels following Antell et al. (2021).

Fossil occurrence data were paired with AOGCM climate data from the midpoint of time bins (e.g., climate was modelled at 12 ka for occurrences binned from 16 ka to 8 ka). Temperature change was calculated as the difference in estimated MAT from one temporal bin to the previous temporal bin for each assemblage. Temperature change was therefore defined locally at the site of the sediment core, providing a best estimate of climate changes experienced by the assemblage. We additionally repeated analysis at a global scale with temperature data derived from paleo-proxies and with AOGCM temperature data derived at depth preferences of individual species (see below).

### Statistical analysis

#### Thermal deviance

We calculated the preferred temperature of each foraminiferal species using an ecological transfer function in the ‘rioja’ R package version 0.9-26 (Juggins, 2020). We used weighted averaging partial least squares (WA-PLS) regression and calibration (Ter Braak, Juggins, Birks, & Van der Voet, 1993) to infer past environmental preferences of planktonic foraminifera assemblages (Imbrie & Kipp, 1971) across MAT at the sea surface. Performance of the WA-PLS transfer function was determined using a leave-one-out cross-validation (see Appendix S1 Fig. S1). WA-PLS requires the development of a calibration dataset, which is then used to model the relationship between assemblages and temperature. We calibrated the WA-PLS function using all planktonic foraminiferal occurrences of the final dataset. We additionally calibrated the WA-PLS function on subsets of the data, focused only on (i) samples from the core top, (ii) samples falling within the interquartile range of all temperature variation throughout the last 700 ka, and (iii) samples falling outside the interquartile range (see Appendix S1 Fig. S2). We calibrated the WA-PLS function on these subsets to test whether calculation of the optimum temperature of each foraminiferal species is strongly dependent on the choice of underlying data. In addition to mean annual SST estimates used in the main analysis, we extracted temperature estimates from each species’ preferred depth. We derived information on each species’ modern depth range from Antell et al. (2021) Table S2, summarised from the literature, and assigned a depth preference as one of the following: surface (40 m in the AOGCM, surface-subsurface (78 m), or subsurface (164 m). WAPLS was then used to re-estimate the preferred temperature of each foraminiferal species.

We define thermal deviance as the difference between the temperature at a given location (as estimated by the AOGCM) and the ‘bio-indicated temperature’, which is the average of the species’ temperature optima within an assemblage, weighted by relative abundances (Fig. 1). Deviance is positive when the water temperature is warmer than the bio-indicated temperature. Previous studies have used the term ‘climatic debt’ for this metric to quantify lags between biotic responses of plants, birds, and butterflies to contemporary climate changes (Bertrand et al., 2016; Devictor et al., 2012; Gaüzère, Princé, & Devictor, 2017). The same metric has been noted as ‘thermal bias’ (Stuart-Smith et al., 2015), ‘community climate lag’ (Blonder et al., 2017), or ‘community-climate mismatch’ (Bonachela, Burrows, & Pinsky, 2021). However, the notation of ‘debt’, ‘lag’, ‘bias’, or ‘mismatch’ might imply that populations occupy suboptimal climate conditions, which is likely not applicable to planktonic organisms. We therefore use the term ‘thermal deviance’ throughout this study.

Thermal deviance calculated by WA-PLS takes the relative abundance of individual species into account. We also estimated thermal deviance based on presence–absence changes in species within assemblages (see Appendix S1 Fig. S3 and S4). To do so, we calculated a species’ temperature preference as the average temperature of the species’ range based on all occurrences of that species across time bins. For a given assemblage, the assemblage temperature was calculated as the average of all species’ temperature indices. Thermal deviance was then calculated as the difference in assemblage bio-indicated temperature and the surface temperature estimate based on the AOGCM. The resulting estimates for the thermal deviance were therefore only based on presence– absence changes of individual species, and do not take the relative abundance of species within assemblages into account.

We modelled the average thermal deviance across sites in a time interval as a function of temperature change using a generalised additive model fitted via the ‘mgcv’ R package version 1.8.41 (Wood, Pya, & Säfken, 2016) and identified change points using the first derivative of the model function. Within latitudinal zones, the trend in thermal deviance was estimated using a linear model; we defined assemblages located between 0° and 30° absolute latitude as low latitude; between 30° and 60° as mid latitude; and above 60° as high latitude. We acknowledge that using absolute latitudes, which merges the Northern and Southern Hemispheres, may introduce biases in the results, particularly for taxa predominantly restricted to one hemisphere, since species with hemispheric preferences may exhibit distinct patterns in turnover dynamics. We therefore repeated our analysis by categorizing latitudinal zones (i.e., mid, high, and low) within hemispheres (see Appendix S1 Fig. S5).

Using the depth-specific estimates of each species, we additionally modelled thermal deviance separately for each depth layer and latitudinal zone using linear models (see Appendix S1 Fig. S6). Assemblages for these models were defined as those species present within a sediment core in a given time bin and given depth habitat.

As a sensitivity analysis, we repeated our modelling steps for thermal deviance using linear mixed effect models in the lme4 R package version 1.1-30 (Bates, Mächler, Bolker, & Walker, 2015). In this approach, we specified temporal bin as a random effect (see Appendix S1 Fig. S7), treating bins as samples of a larger population to account for variability and differences across bins. Additionally, we fitted a first-order autoregressive model to estimate latitudinal trends in thermal deviance to account for temporal non-independence (see Appendix Table S3). We further tested whether the substantially increased number of samples towards the recent affected our results by repeating our analysis on a subset of data that omitted samples from the most recent time bin, containing the highest number of samples (1,901 of 2,657 assemblages; see Appendix S1 Fig. S7, S8 and S9).

#### Compositional turnover

We were interested in the relationship between degree of compositional turnover and climate change through time and space. We therefore calculated compositional turnover at each latitudinal zone across consecutive time steps. We grouped all assemblages occurring in the same temporal interval and latitudinal zone together, and calculated turnover against all assemblages of the same latitudinal zone of the previous bin. For this, we calculated dissimilarity indices with the ‘vegan’ R package version 2.6.2 (Oksanen et al., 2020) based on the Chi-squared coefficient (Prentice, 1980), which can be considered a more robust method for relative abundance assemblage data than the classical Bay-Curtis coefficient (Mottl et al., 2021). We then modelled turnover per site and temporal bin as a function of per-latitudinal zone temperature change using linear models. We repeated this analysis by using the more traditional Bray-Curtis dissimilarity index instead of Chi-squared coefficients (see Appendix Table S4).

#### Species richness

We additionally counted the number of species per latitudinal zone for each temporal bin and modelled this proxy of species richness as a function of temperature change using linear models.

#### Extirpation probability

To test whether high thermal deviances correspond to local extinctions for individual species within assemblages, we quantified the average extirpation probability for each latitudinal zone under instances of climate warming and climate cooling. For this analysis, we counted the number of species that were present in a latitudinal zone in temporal bin *i* but not in the subsequent temporal bin *i+1*. We additionally noted whether the temperature change was above zero (i.e., climate warming) or below zero (i.e., climate cooling) from temporal bin *i* to temporal bin *i+1* for the focal latitudinal zone. The extirpation probability was then modelled in a time-continuous logistic regression for all temporal bins and summarised per climate scenario (i.e., warming and cooling).

#### Temperature sensitivity analysis

We calculated thermal deviance for planktonic foraminifera assemblages based on the mean annual surface temperature estimate of the AOGCM for the main-text results. As both the assemblage bio-indicated temperature and the temperature reconstructions are based on AOGCMs, however, we additionally tested whether the same patterns were observed on a global level when using a recent compilation of paleo-proxy data for temperature reconstructions instead (Friedrich & Timmermann, 2020). The paleo-proxy data spanned the last 125 ka and was converted to the same resolution as our assemblage data by averaging proxy values for the temporal bins of the assemblage data (i.e., upscaling or aggregation). We tested whether taking a point estimate for the temperature reconstruction biased our results by iteratively sampling the age estimates of the paleo-proxy data from the entire distribution, including the temperature estimates for the mean age ± 4 ka (see Appendix S1 Fig. S10), which corresponds to the temporal uncertainty of our fossil data.

## Results

### Global thermal deviance

Planktonic foraminiferal assemblages generally tracked temperature changes over the past 700,000 years. However, assemblages showed little compositional change when temperature changes were below ca. 0.3°C, which resulted in thermal deviances between the actual temperature at the site and the bio-indicated temperature of the assemblage (Fig. 2b). Under larger temperature changes (> 0.3°C), thermal deviances remained nearly constant.

**Figure 2:**
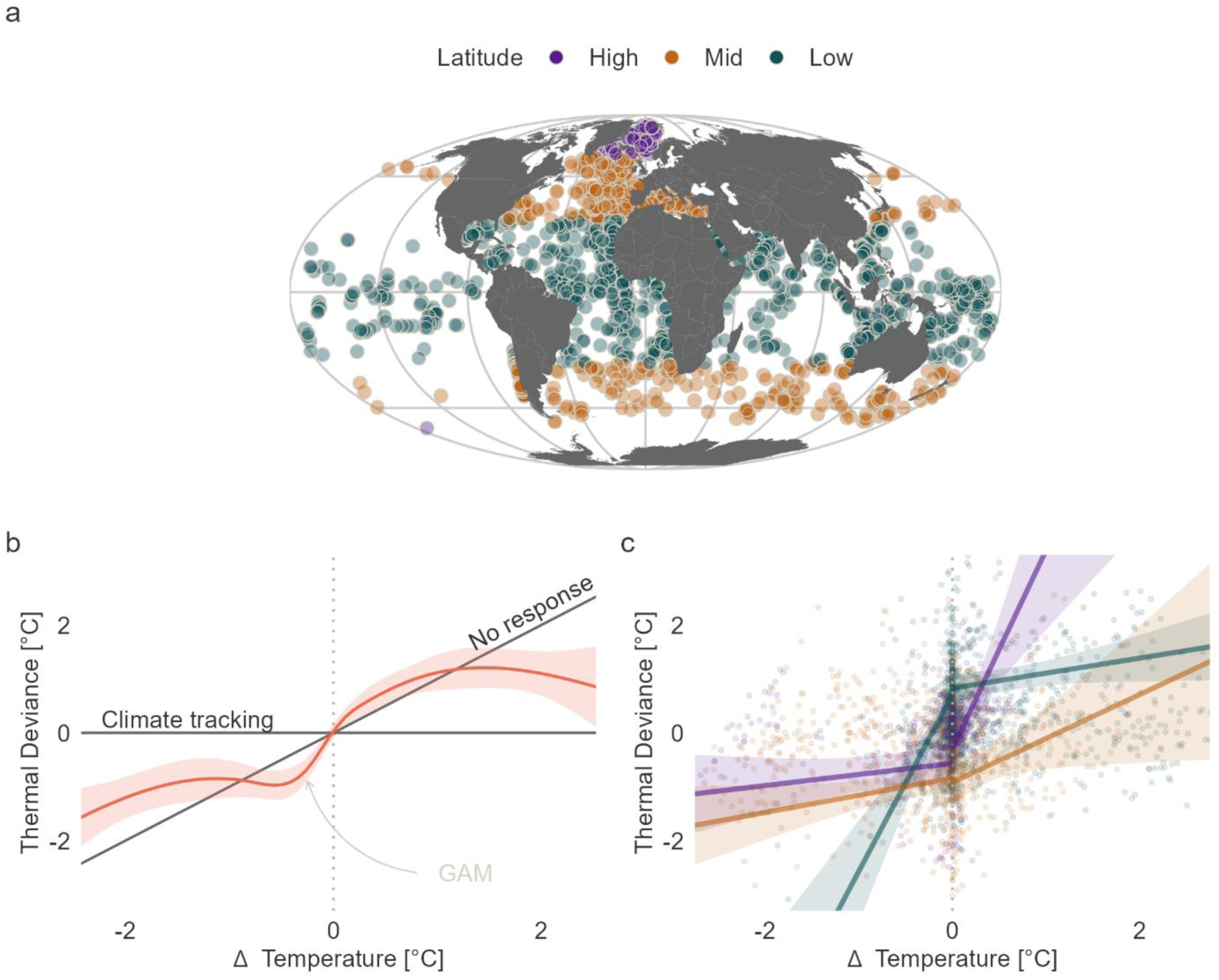
(a) Global distribution of foraminifera assemblages used in this work, coloured by latitudinal zones, 0-30°, 30-60°, 60-90° absolute latitude. (b) A generalised additive model (GAM) predicts thermal deviance for each assemblage as a function of estimated local temperature change from the previous time step. The deviation is itself the difference between coeval local temperature estimated from Earth systems models (AOGCM) vs. the temperature expected based on assemblage composition and abundance. The horizontal line shows a one-to-one relationship between AOGCM temperature and bio-indicated temperature (e.g., 1°C warming corresponds to a bio-indicated temperature that is 1°C warmer, resulting in no thermal deviance). The ‘no response’ line shows a constant bio-indicated temperature regardless of AOGCM temperature change. (c) Latitudinal patterns of thermal deviance as a function of temperature change. The coloured areas depict the 95% confidence interval of the focal trend based on linear regression models. A positive deviation corresponds to bio-indicated temperatures adjusting to be cooler than expected from Earth systems model temperature estimates, and a negative deviation to warmer bio-indicated temperatures than expected.

Our results remain consistent when using global temperature estimated from paleo-proxies instead of modelled AOGCM temperature estimates (Appendix S1 Fig. S10), indicating that relationships described here are not an artefact of using AOGCM output for both the modelled temperature and the thermal deviance estimates.

### Thermal deviance across latitudes

The response of planktonic foraminiferal assemblages to temperature change through time was spatially heterogeneous (Fig. 2c, Appendix S1 Table S3). Assemblages at high latitudes showed moderate thermal deviance during climate cooling but accumulated substantial thermal deviance with warming (Fig. 2c). With every 1°C warming, average bio-indicated temperatures of high latitude assemblages were 3.5°C lower than the actual temperature at the site (95% CI [2.1°C, 5°C]; Appendix S1 Table S3). Conversely, assemblages at low latitudes exhibited higher thermal deviances under climatic cooling and smaller deviances when temperatures increased. With every 1°C cooling, average temperature preferences of low latitude assemblages were 2.7°C warmer than the site temperature (95% CI [1.6°C, 3.8°C]; Appendix S1 Table S3). Assemblages at mid latitudes closely tracked temperature cooling and showed a modest increase in thermal deviance under temperature warming.

Using various data subsets for estimation of the ecological transfer function (see Appendix S1 Fig. S2) and omitting data from time bins that showed indications of higher or lower sampling intensity (see Appendix S1 Fig. S4) produced the same basic results of spatially heterogenous responses to temperature change. Similarly, using a mixed effect model with a random effect on temporal bins resulted in the same patterns (Appendix S1 Fig. S7), in accordance with autoregressive models accounting for temporal independence (Appendix S1 Table S3). The estimates for the overall direction of the relationship between thermal deviance and temperature change fall within the 95% confidence intervals of the original data apart from low latitude assemblages under climate warming, which show a slightly decreased thermal deviance with increasing temperature change. The same applies when defining latitudinal zones within hemispheres (Appendix S1 Fig. S5).

Using temperature estimates derived at each species’ preferred depth layer to model thermal deviance resulted in the same trends as for surface temperature, apart from mid latitude assemblages during climate cooling (see Appendix S1 Fig. S6). High latitude assemblages showed large increases in thermal deviance during climate warming, and low latitude assemblages during cooling. Mid latitude assemblages, however, showed larger increases in thermal deviance during cooling as estimated by surface temperature.

Calculating thermal deviance on occurrences instead of relative abundance of individual foraminifera species revealed the same trends but generally higher magnitudes of thermal deviance, at both the global scale (see Appendix S1 Fig. S3) and within latitudinal bands (see Appendix S1 Fig. S4).

### Assemblage composition

Compositional turnover increased for mid latitude assemblages with increasing magnitude of temperature change under both warming and cooling scenarios (Fig. 3a, Appendix S1 Table S4). This was not the case for high and low latitude assemblages. High latitude assemblages showed an increase in compositional turnover during climate cooling but not during climate warming, where compositional turnover generally decreased (Fig. 2c, Appendix S1 Table S4). In contrast, low latitude assemblages showed slightly increasing to constant compositional turnover with climate warming and constant compositional turnover with climate cooling. We obtained the same results using the more traditional Bray-Curtis dissimilarity index instead of Chi-squared coefficients (see Appendix S1 Table S4).

**Figure 3:**
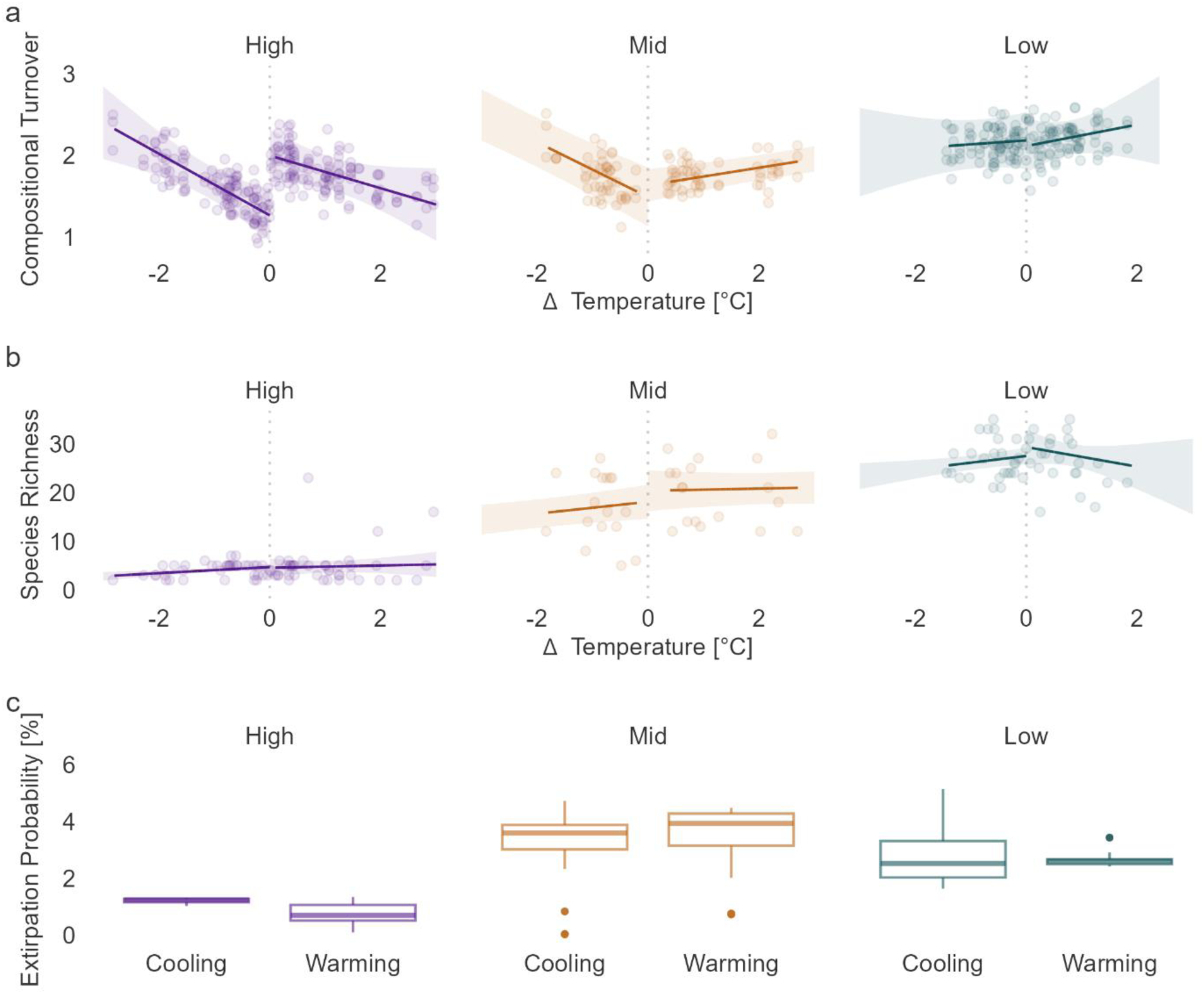
Changes in assemblage composition as a response to temperature changes. (a) Compositional turnover for each latitudinal zone based on Chi-squared dissimilarity indices for each assemblage. (b) The number of species within each latitudinal zone. (c) Extirpation probability as modelled by the binomial signal of remaining species against species that experienced extirpation in an assemblage through time (see Methods). The bold line in (a) and (b) depicts the mean trend across data points, and the shaded area the corresponding 95% confidence interval.

Species richness was highest in low latitudinal zones and lowest in high latitudinal zones (Fig. 3b). Species richness was generally constant during climatic changes (see Appendix S1 Table S5).

We found no substantial differences in extirpation probabilities between cooling and warming scenarios for all latitudinal zones (Fig. 3c). Average extirpation probability was highest in mid latitude assemblages and lowest in high latitude assemblages (see Appendix S1 Table S5).

## Discussion

Our findings suggest that assemblages of planktonic foraminifera track temperature changes, which is congruent with existing literature on marine microfossils on global (Strack et al., 2022; Yasuhara, Hunt, Dowsett, Robinson, & Stoll, 2012; Yasuhara, Tittensor, Hillebrand, & Worm, 2017) and local (Bond et al., 1997; Field, Baumgartner, Charles, Ferreira-Bartrina, & Ohman, 2006; Hüls & Zahn, 2000) spatial scales. However, assemblages tolerate modest temperature changes (< 0.3°C) with only little change in composition. Additionally, under climate warming, more cold-adapted species were found in polar assemblages than predicted by a linear, one-to-one relationship between water temperature and bio-indicated temperature. Similarly, we observed a disproportionate abundance of warm-adapted species in equatorial assemblages under a cooling scenario. These results are consistent with the persistence and survival of species with the highest or lowest temperature preferences, regardless of the magnitude of climate change over glacial–interglacial cycles. At mid latitudes, the bio-indicated temperatures of assemblages mirrored the corresponding temperature change more closely.

These latitudinally-heterogeneous responses of assemblages to temperature change can be linked to observed patterns in compositional turnover (Fig. 3a). At a given site undergoing warming, more warm-adapted species are expected to move to the site (or increase their in situ abundance) and more cold-adapted species are expected to go locally extinct (or decrease their in situ relative abundance), and vice versa for cooling (e.g., Chen et al., 2011). Both processes contribute to turnover. In accordance with these expectations, we observed an increase in compositional turnover in mid and low latitude assemblages with warming (Fig. 3a). In contrast, compositional turnover tends to decrease or stay constant with magnitude of warming in high latitude assemblages, increasing the observed thermal deviance of high latitude assemblages under climate warming (Fig. 2c). Similarly, while compositional turnover increases during climate cooling for high and mid latitudinal assemblages as expected, turnover is constant for low latitude assemblages, resulting in an increasing thermal deviance for the latter.

Our results highlight the resilience and inertia of planktonic foraminiferal assemblages to climate change. Modest temperature changes are likely to have remained within the thermal tolerance limits of many species within an assemblage (see Supporting Information Appendix S1 Table S3), such that species did not experience pressure to shift their distributions or abundance, resulting in a high compositional stability at the assemblage level. We observed this for temperature changes below 0.3°C per 8 ka on a global level (Fig. 2b). Under larger temperature changes (> 0.3°C), the persistence of high latitude assemblages during climate warming and low latitude assemblages during climate cooling (Fig. 2c) did not lead to increased extirpation risk (Fig. 3c). This can be attributed to several potential, non-mutually exclusive mechanisms. First, thermal conditions might always remain within the thermal tolerance limits of the coldest- and warmest-adapted species. These species might be able to survive wide deviations from their thermal optimum, for instance by remodelling their physiology in situ (Seebacher, White, & Franklin, 2015). Alternatively, thermal heterogeneity at small spatial scales and/or across a depth gradient may provide microhabitats suitable for species at current locations, even as regional average temperatures begin to change (Fuller et al., 2010; Kretschmer, Jonkers, Kucera, & Schulz, 2018). While autotrophic foraminifera may be more constrained in their suitable vertical distribution by light availability, all planktonic species have some leeway to vary their depth distribution and thereby moderate exposure to large-scale changes in environmental conditions.

However, high resilience may only partially explain the decreasing turnover of high-latitude assemblages during climate warming (Fig. 3a), which is in contrast to expectations of general turnover trends (e.g., Chen et al., 2011). While interspecific competition is unlikely to impede species immigration (Rillo et al., 2019), factors such as sea ice cover, seasonality, ocean chemistry, and light-availability may contribute to the observed pattern (Zamelczyk et al., 2021). We therefore emphasize that the relationship between thermal deviance, extirpation risk, and turnover may depend on numerous ecological and environmental factors.

An alternative explanation for the observed latitudinal signal in thermal deviance could be that modelled temperature estimates from the AOGCM are directionally biased, such that thermal deviances would be simply emerging erroneously from the climate model output. Prior studies suggest such errors in paleoclimate models may be localized, for example to the North Atlantic during glacial intervals (Jonkers et al., 2023). Thus, although imprecision in estimated sea temperature exists in our study, it is improbable it explains the symmetric observations at equatorial and polar latitudes, as well as the consistency of results under various robustness tests (see Material and Methods and Results section). The climate model output used here also generally aligns with temperature estimates derived from paleo-proxy data (Braconnot et al., 2012; Tierney et al., 2020).

Our results further highlight the importance of *within-community* relative abundance changes for explaining species’ responses to temperature changes. We found higher thermal deviances when relative abundance changes were omitted, indicating species responded to warming or cooling through substantial abundance changes within an assemblage. Novel plankton communities, such as those described since the last ice age (Strack et al., 2022), could therefore result from both habitat tracking and *in situ* abundance changes, and both mechanisms need to be incorporated when assessing the biotic response to temperature change.

We emphasise that our results only estimate compositional changes of assemblages as a function of temperature. Although temperature is likely the single most important explanatory variable structuring the geographic distributions of planktonic foraminifera (Fenton et al., 2016; Yasuhara, Wei, et al., 2020), other environmental parameters can similarly affect species turnover, particularly in warmer waters such as above 25° (Rillo, Woolley, & Hillebrand, 2022). The high thermal deviance of low-latitude assemblages under cooling reported here may therefore potentially result from a decoupling of temperature and species turnover dynamics in warm ecosystems due to the increased importance of other environmental factors such as net primary productivity or water depth (Rillo et al., 2022). Using the results reported here to estimate future compositional change of planktonic assemblages, e.g. under anthropogenic climate change, requires careful consideration of potential interacting environmental parameters and temporal scales. Nevertheless, our results can be taken to suggest that planktonic foraminifera, and perhaps other planktonic organisms, are generally able to closely track temperature changes, albeit with spatially distinct responses of high and low latitude assemblages.

## Data Accessibility Statement

All data and code files are publicly available at https://github.com/Ischi94/climatic_debt. All analyses were carried out in R v.4.1.2 (R Core Team, 2023). We used the ‘tidyverse’ collection of R packages (Wickham et al., 2019) to transform and visualise data.

## Acknowledgements

This work was supported by the Deutsche Forschungsgemeinschaft (KI 806/16-1, AB 109/11-1 and STE 2360/2-1/ FOR 2332: Temperature-related stressors as a unifying principle in ancient extinctions).

