## Supporting Information Appendix S1 for "Spatially heterogeneous responses of planktonic foraminifera assemblages over 700,000 years of climate change"

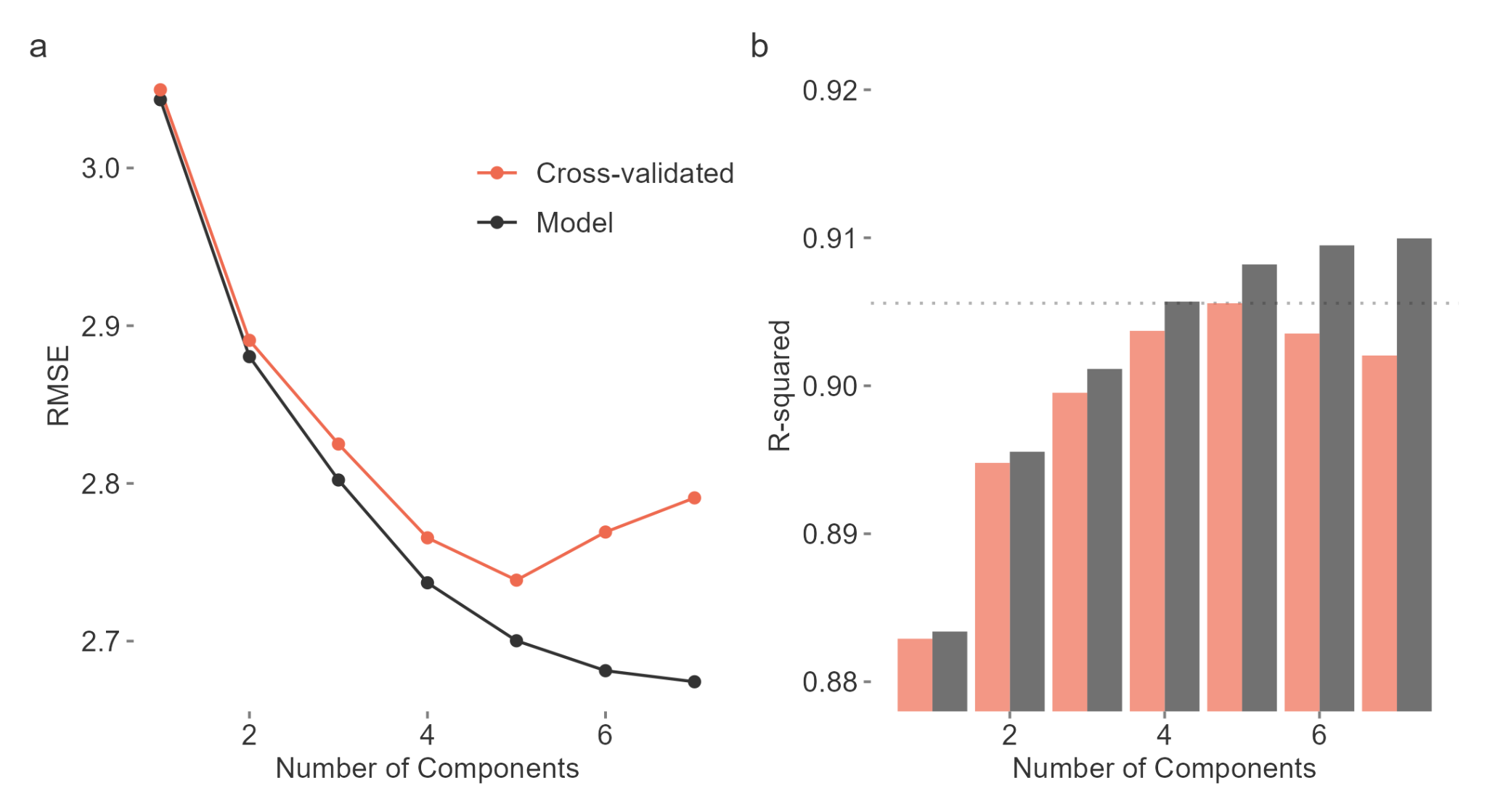

Fig. S1: Weighted averaging partial least squares (WA-PLS) regression was used to estimate the relationship between assemblage composition of planktonic foraminifera and ambient temperature. (a) The performance of the model was assessed on the training subset (black line and points) and by leave-one-out cross-validation (orange line and points) via the root mean square error (RMSE). The appropriate number of components to use in the subsequent analysis was chosen by minimising the cross-validated RMSE, which resulted in 5 components. (b) The R^2^ value for the WA-PLS regression based on cross-validation (orange) and test subset (black) as a function of the number of components.

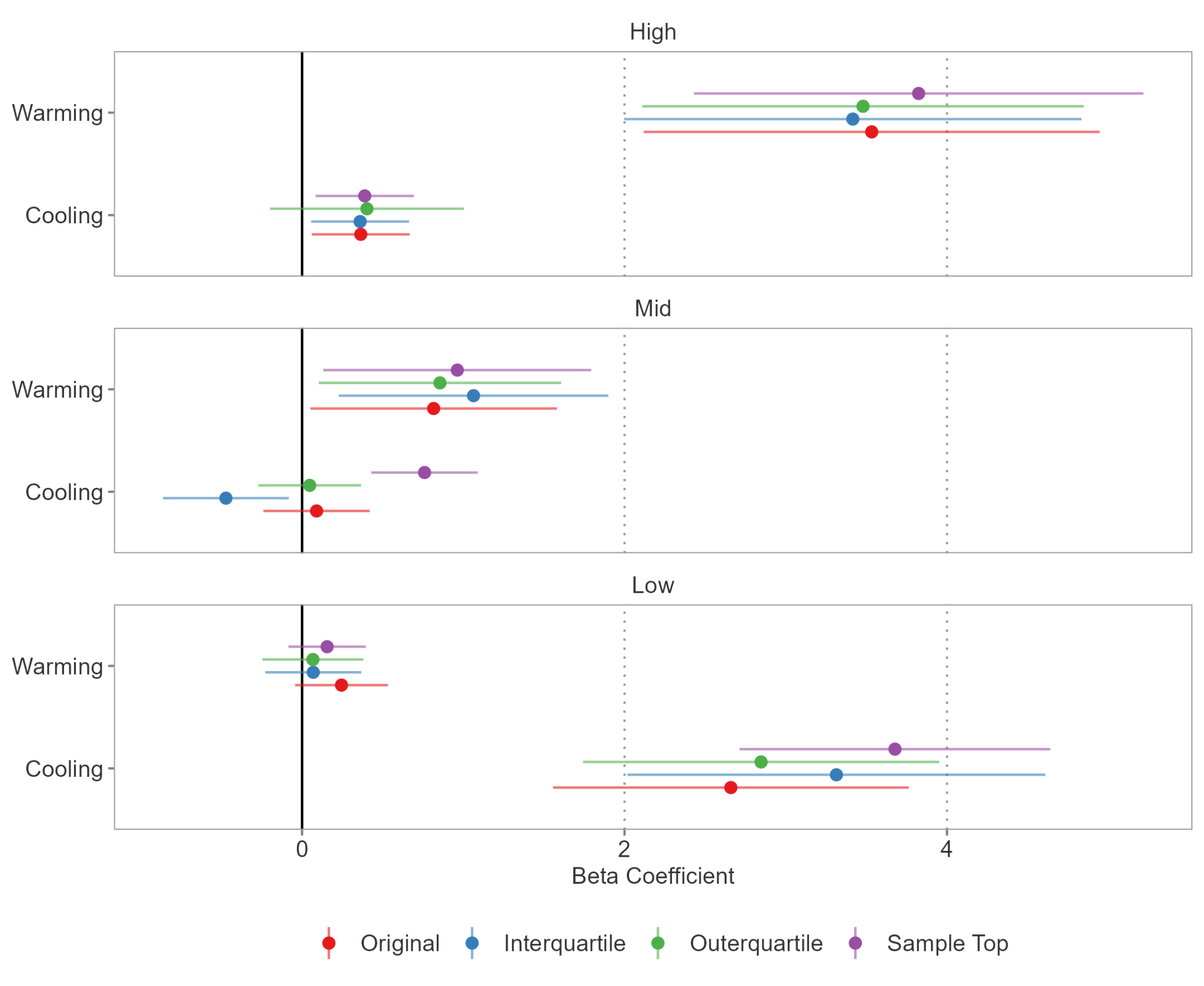

Fig. S2: Results for the global thermal deviance of all foraminifera assemblages as a function of temperature change (Beta Coefficient and 95% Confidence Interval) as estimated with various datasets for the ecological transfer function. The original estimate is based on the whole dataset (red). The blue points and lines indicate estimates based on data points falling within the interquartile range of all temperature variation throughout the last 700 ka, whereas green lines and points show the estimates based on data points falling outside of this range. The purple estimates are based on samples from the sample top only.

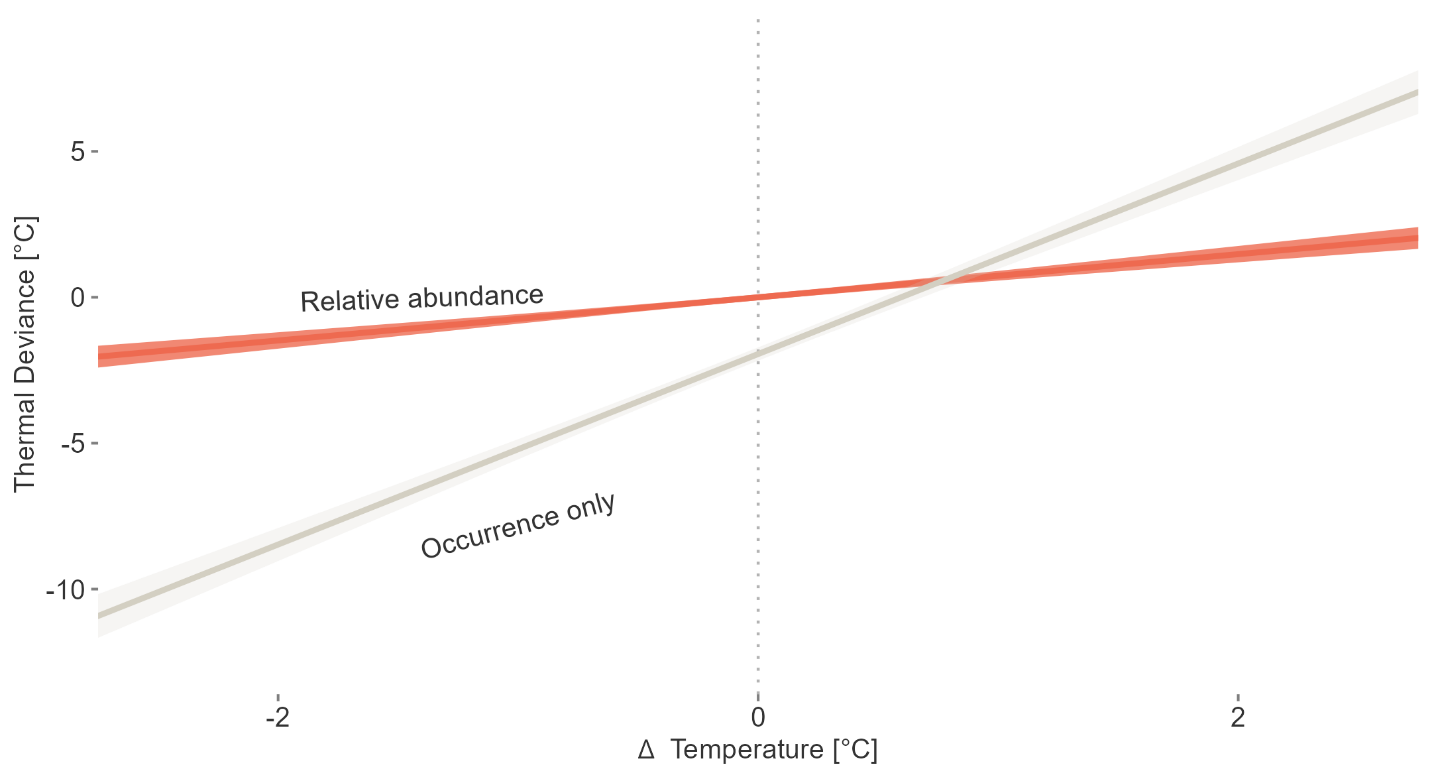

Fig. S3: The global thermal deviance of all foraminifera assemblages as a function of temperature change, based on the preferred temperatures of assemblages estimated by including the relative abundance of individual species (orange line), or by occurrences only (grey line). The colored area depicts the focal 95% confidence interval of the regression slope.

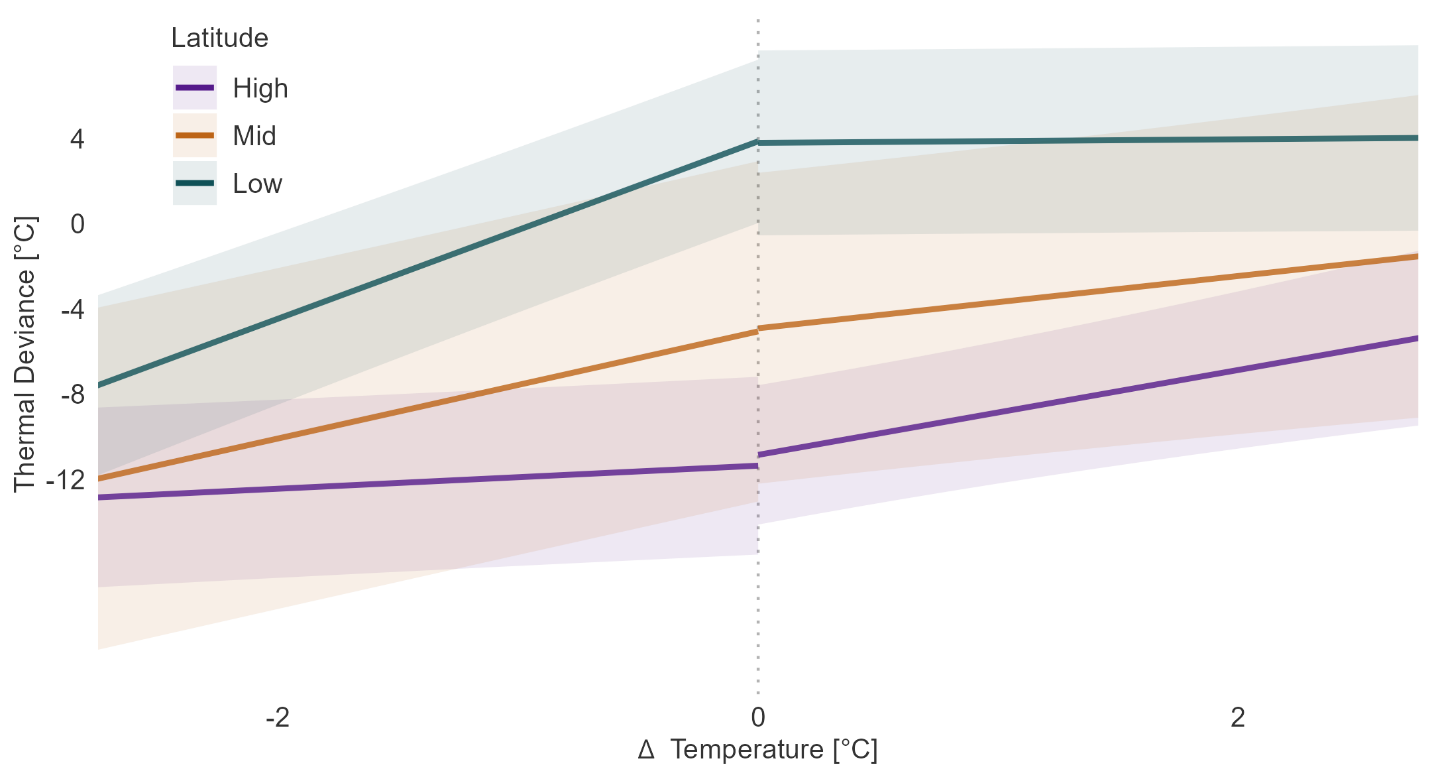

Fig. S4: The thermal deviance of foraminifera assemblages per latitudinal zone as a function of temperature change, based on the bio-indicated temperatures of assemblages estimated by occurrences only. The colored areas depict the 95% confidence interval of the focal regression slope.

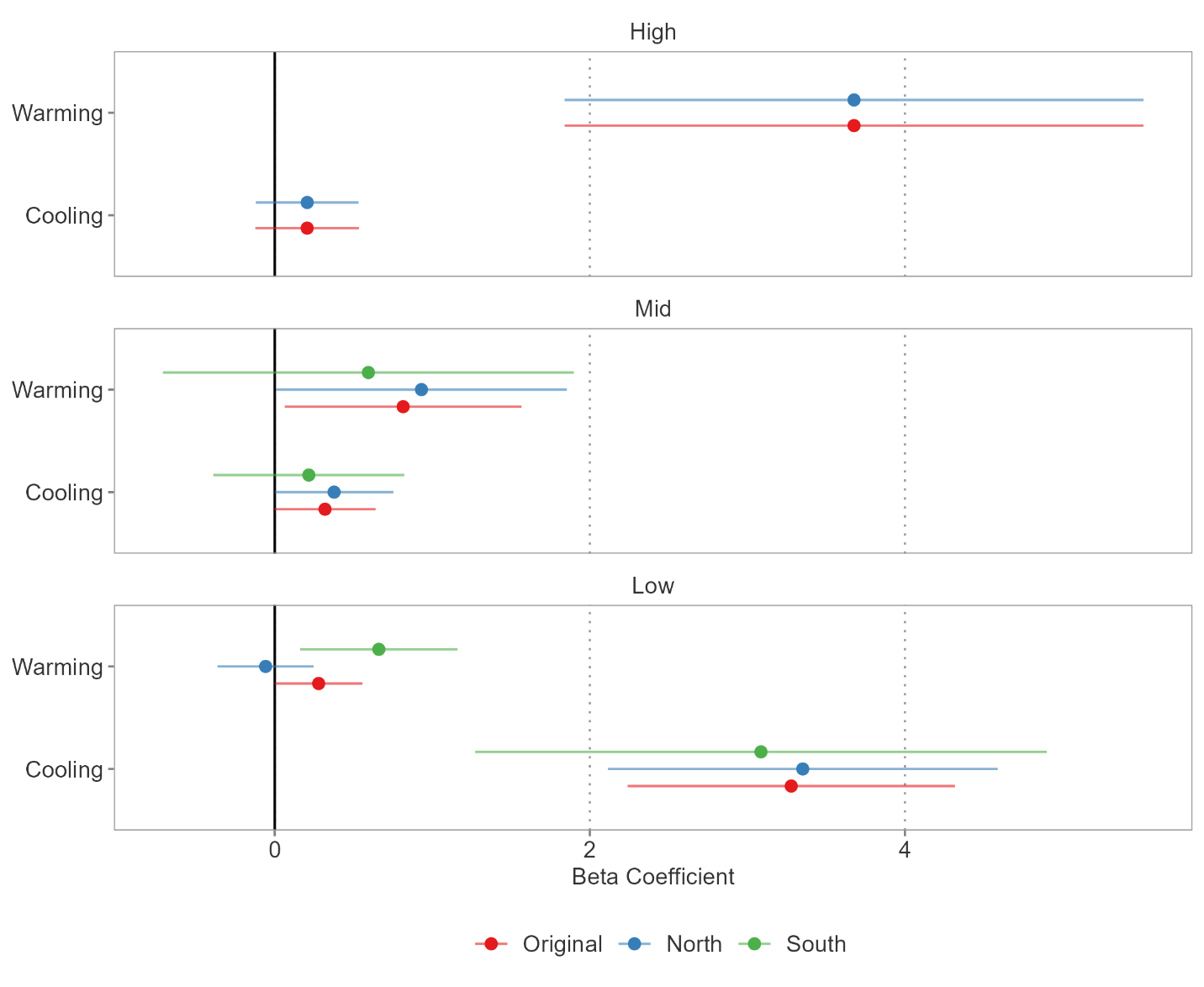

Fig. S5: Results for the global thermal deviance of all foraminifera assemblages as a function of temperature change (Beta Coefficient and 95% Confidence Interval) as estimated by latitudinal zones defined by absolute latitudes (merging the Northern and Southern Hemispheres, Original) and by latitudinal zones within hemispheres. The red estimates are reported throughout the text. Blue estimates are based on the Northern Hemisphere and green estimates on the Southern Hemisphere.

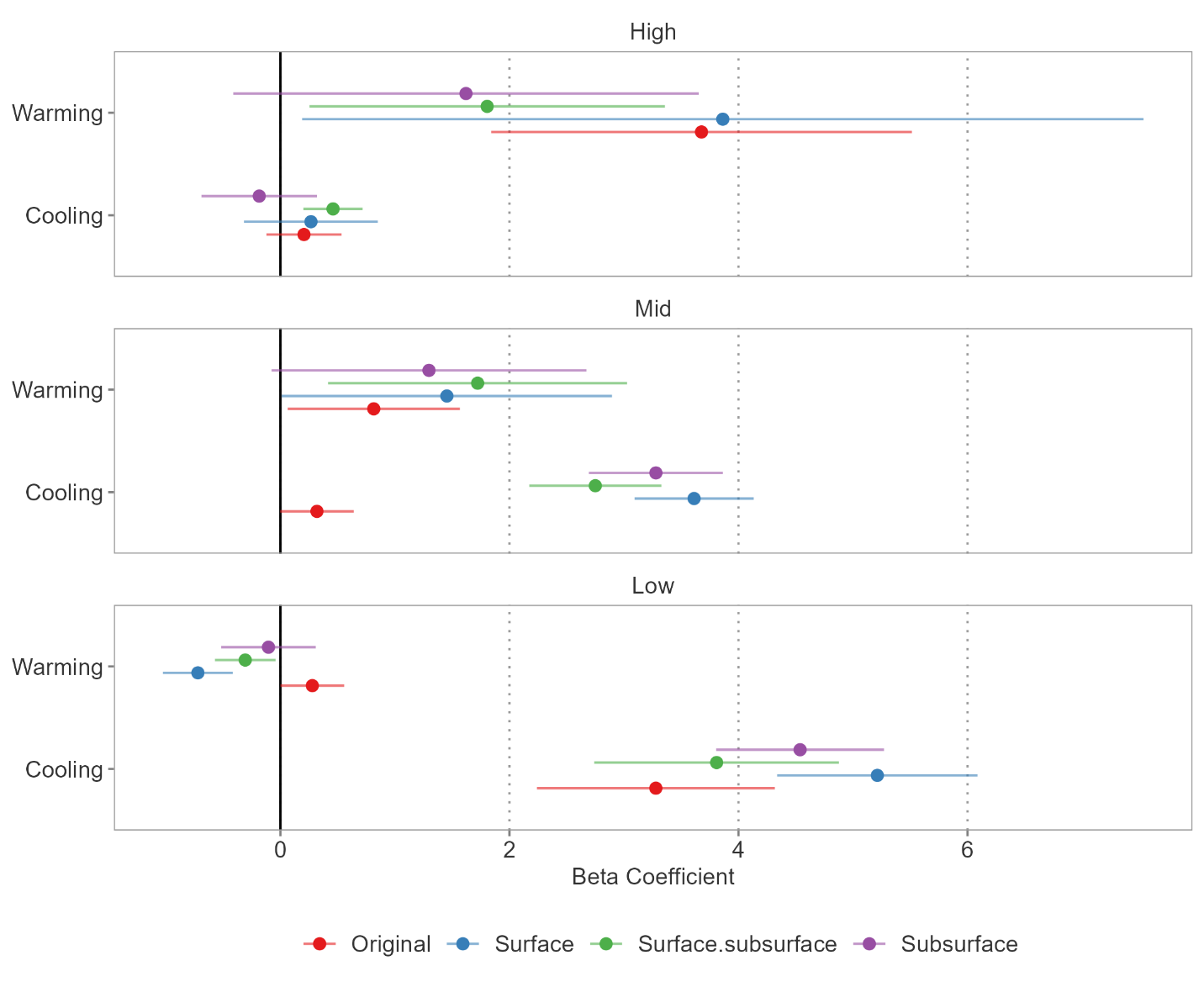

Fig. S6: Results for the global thermal deviance of all foraminifera assemblages as a function of temperature change (Beta Coefficient and 95% Confidence Interval) as estimated with surface temperature estimates (Original) and depth habitat preferences of individual species. The red estimates are reported throughout the text and are based on the full data set and fixed effects regression models. Blue estimates are based on the AOGCM surface layer (40m), green estimates on the surface-subsurface layer (78m) and purple estimate on the subsurface layer (164m).

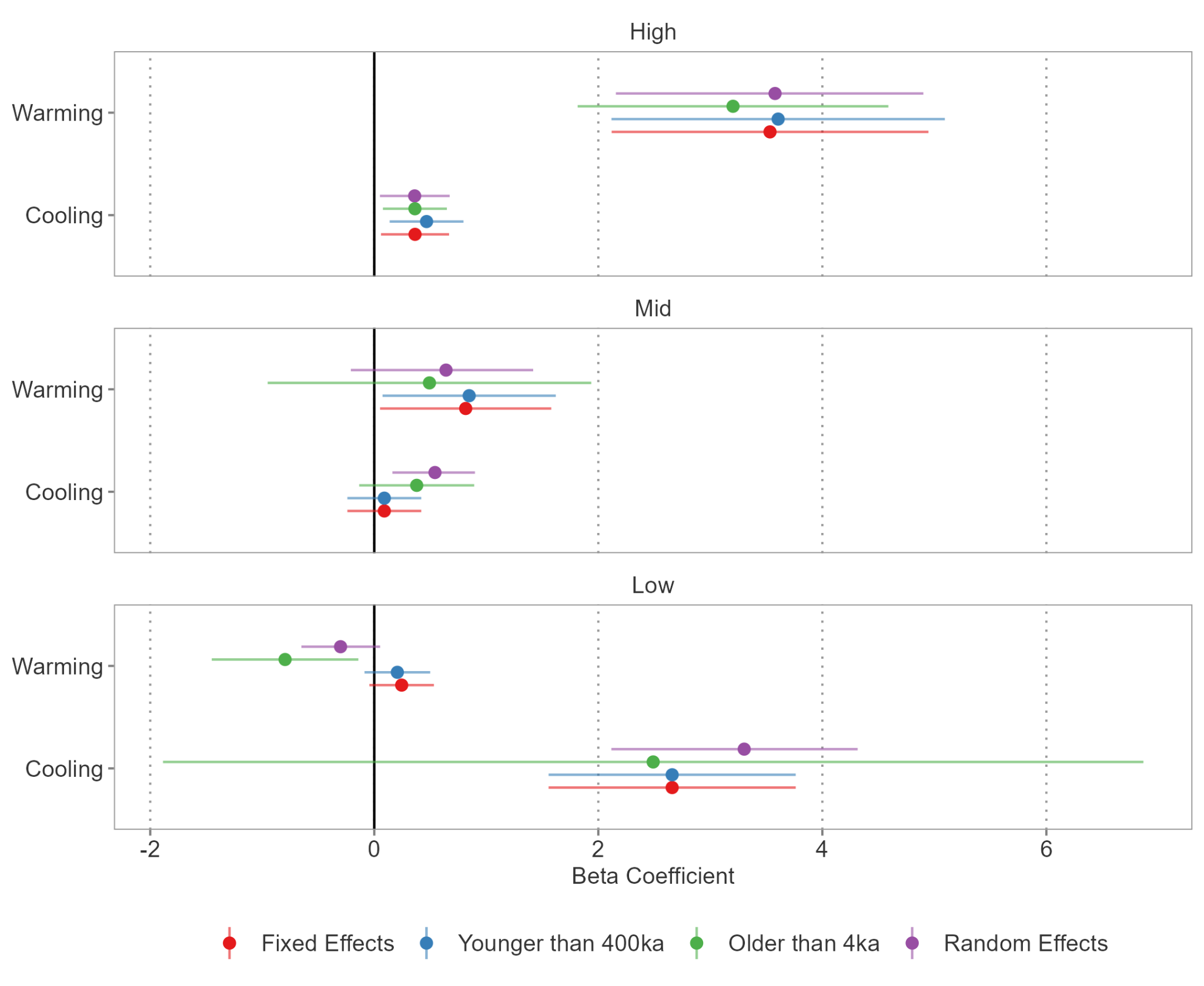

Fig. S7: Results for the global thermal deviance of all foraminifera assemblages as a function of temperature change (Beta Coefficient and 95% Confidence Interval) as estimated with various data subsets and models. The red estimates are reported throughout the text and are based on the full data set and fixed effects regression models. Blue estimates are based on a data subset where all observations older than 400 ka were omitted, while green estimates are based on a subset where the youngest bin (4 ka) was omitted. The purple estimates are based on the full data set and random effects models, where a random effect was put on bins in order to account for non-independence of observations through time.

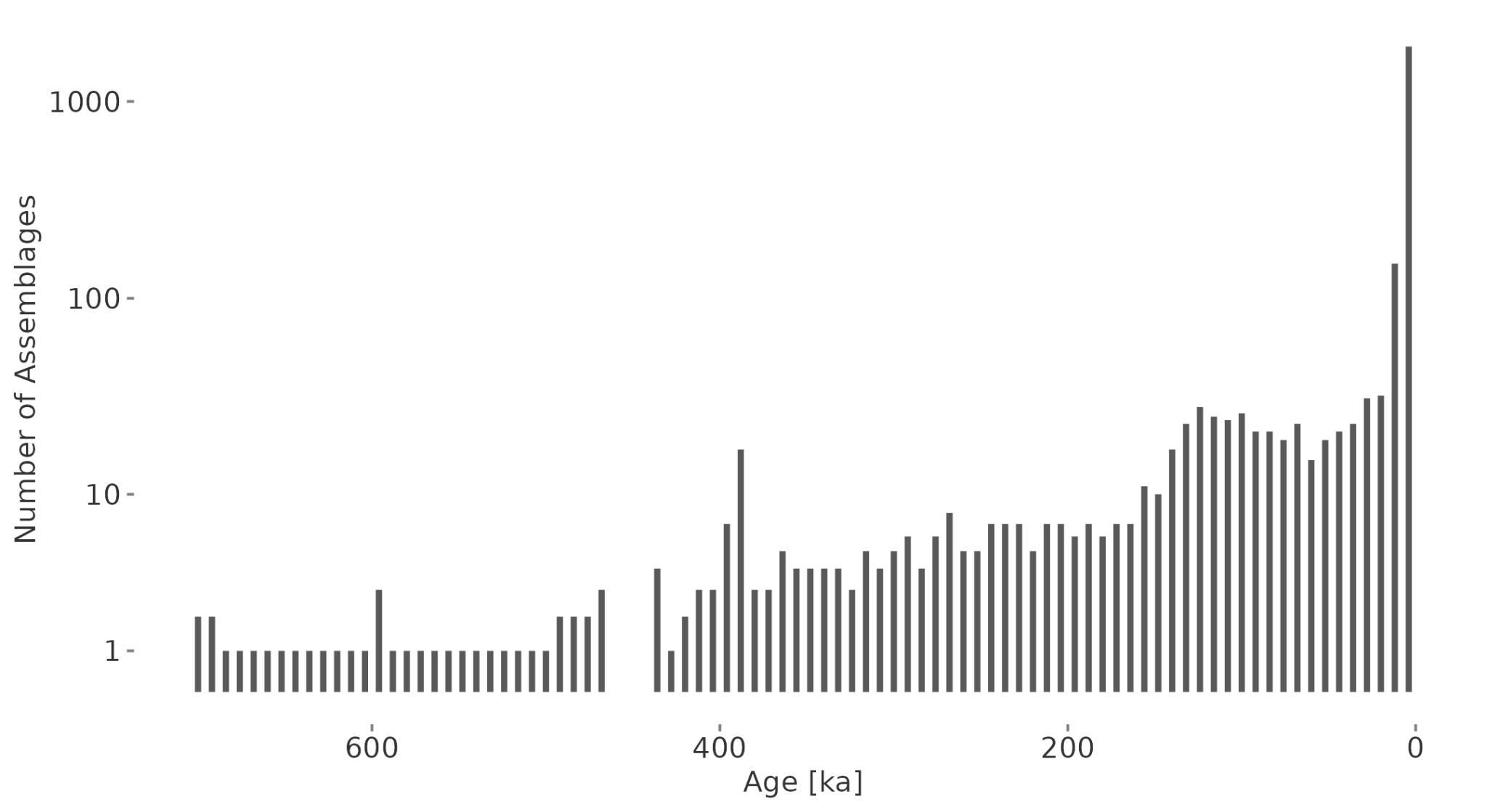

Fig. S8: The number of planktonic foraminifera assemblages through time. As planktonic foraminifera assemblages were unsampled in some time bins between 460 ka and 444 ka, the resulting global bin-to-bin thermal deviance around this interval can be considered unreliable. Note that the number of assemblages is plotted on a logarithmic scale.

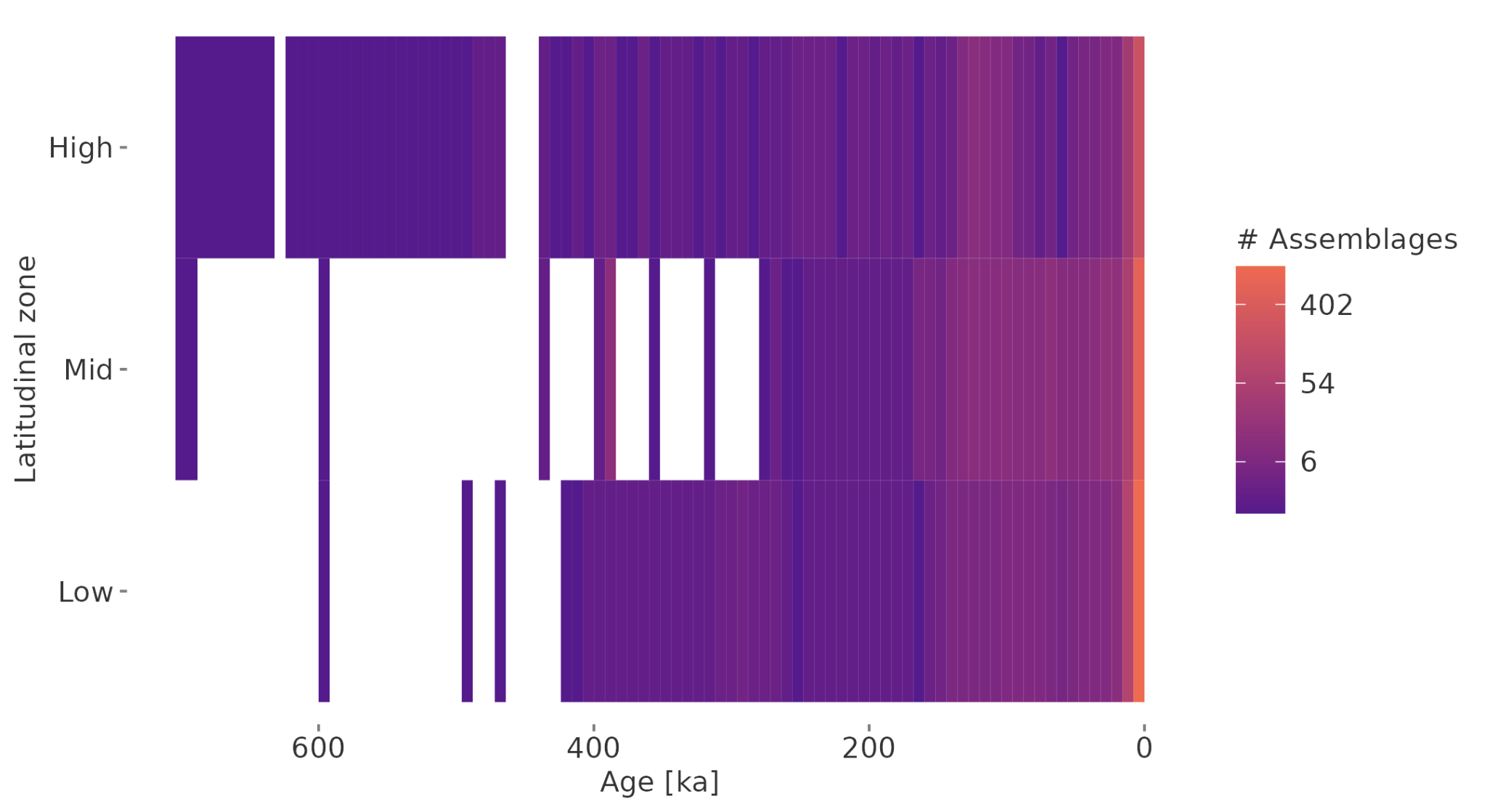

Fig. S9: The number of planktonic foraminifera assemblages through time and space. As planktonic foraminifera assemblages were unsampled in some time bins between 460 ka and 444 ka, the resulting global bin-to-bin thermal deviance around this interval can be considered unreliable. Note that the number of assemblages is plotted on a logarithmic scale.

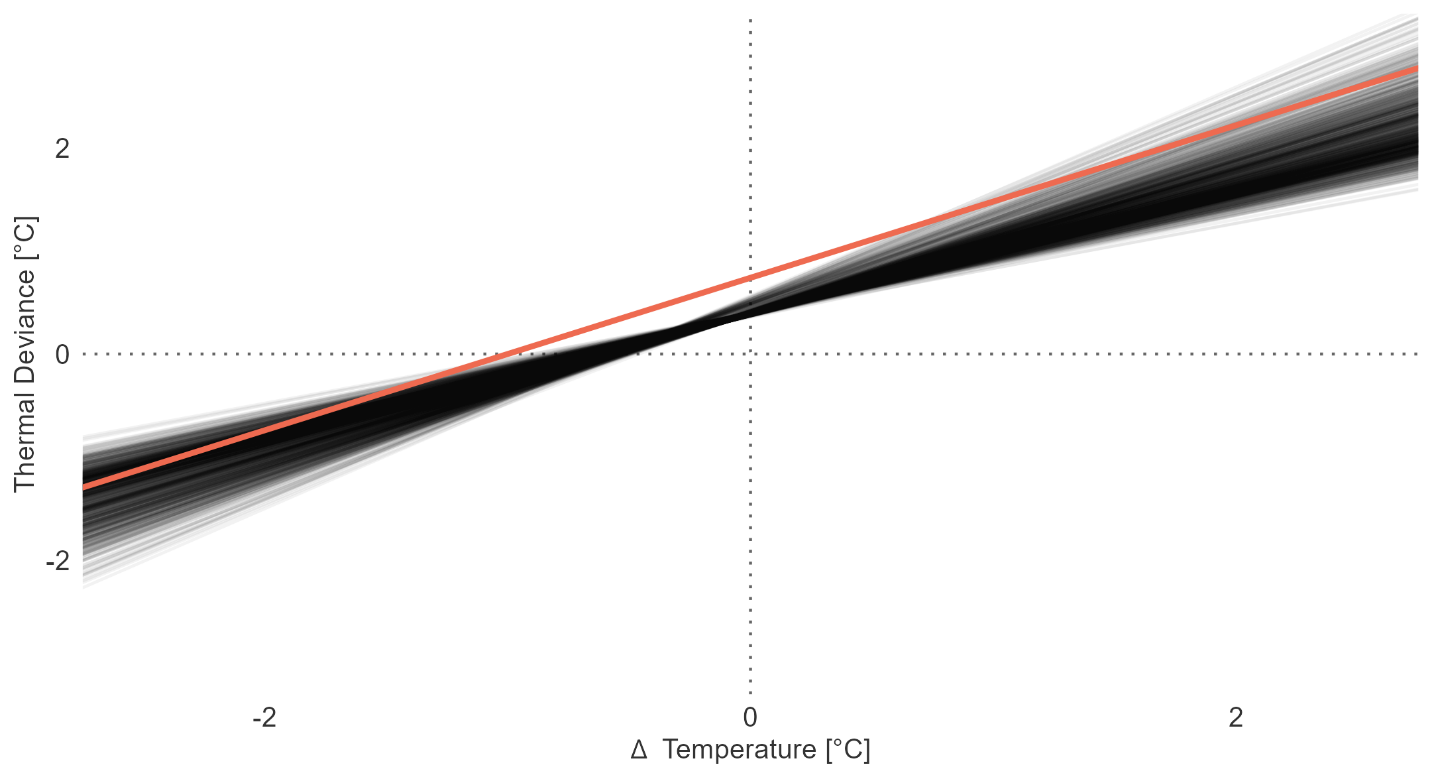

Fig. S10: The linear trend in thermal deviance as a function of temperature change. The colored trend line shows the estimated trend based on AOGCM output for temperature estimates, and the grey lines shows 1000 bootstrapped trend lines based on paleo-proxy data for temperature estimates. As paleo-proxy data was available at a higher resolution (1 ka) than the compositional data (8 ka), we iteratively sampled paleo-proxy estimates around ±4 ka of the age of each temporal bin of the compositional data.

Table S1: Number of occurrences per species for the cleaned dataset for the whole time period (total) and latitudinal zones, 0-30, 30-60, 60-90 degrees absolute latitude. The dataset contains 55 species with 38,441 total occurrences.

| **Species** | **Total** | **High** | **Mid** | **Low** |
| --- | --- | --- | --- | --- |
| Beella digitata | 1,108 | 4 | 339 | 765 |
| Beella megastoma | 12 | 12 |  |  |
| Beella praedigitata | 7 |  | 4 | 3 |
| Candeina nitida | 270 |  | 19 | 251 |
| Globigerina bulloides | 2,272 | 221 | 909 | 1,142 |
| Globigerina falconensis | 1,487 | 31 | 582 | 874 |
| Globigerina umbilicata | 8 |  | 8 |  |
| Globigerinella adamsi | 66 |  |  | 66 |
| Globigerinella calida | 1,447 | 1 | 412 | 1,034 |
| Globigerinella siphonifera | 1,787 | 5 | 524 | 1,258 |
| Globigerinita glutinata | 2,235 | 165 | 820 | 1,250 |
| Globigerinita minuta | 7 |  |  | 7 |
| Globigerinita uvula | 80 | 26 | 37 | 17 |
| Globigerinoides conglobatus | 1,077 |  | 220 | 857 |
| Globigerinoides elongatus | 5 |  | 1 | 4 |
| Globigerinoides ruber | 1,889 | 8 | 606 | 1,275 |
| Globoconella inflata | 1,574 | 112 | 842 | 620 |
| Globoconella puncticulata | 3 |  | 3 |  |
| Globoquadrina conglomerata | 334 | 2 | 14 | 318 |
| Globorotalia anfracta | 47 |  | 1 | 46 |
| Globorotalia flexuosa | 5 |  |  | 5 |
| Globorotalia tumida | 979 | 3 | 65 | 911 |
| Globorotalia ungulata | 9 |  |  | 9 |
| Globorotaloides hexagonus | 338 |  | 35 | 303 |
| Globoturborotalita rubescens | 1,067 | 1 | 264 | 802 |
| Globoturborotalita tenella | 1,175 | 1 | 279 | 895 |
| Hastigerina pelagica | 43 |  | 1 | 42 |
| Hastigerinella digitata | 29 |  | 12 | 17 |
| Hirsutella hirsuta | 825 | 12 | 454 | 359 |
| Hirsutella scitula | 1,350 | 35 | 573 | 742 |
| Hirsutella theyeri | 128 |  |  | 128 |
| Menardella menardii | 1,274 |  | 161 | 1,113 |
| Neogloboquadrina dutertrei | 1,738 | 37 | 487 | 1,214 |
| Neogloboquadrina humerosa | 1 |  | 1 |  |
| Neogloboquadrina incompta | 2,017 | 342 | 851 | 824 |
| Neogloboquadrina pachyderma | 1,455 | 358 | 717 | 380 |
| Orbulina suturalis | 10 |  | 10 |  |
| Orbulina universa | 1,709 | 28 | 631 | 1,050 |
| Pulleniatina finalis | 1 |  |  | 1 |
| Pulleniatina obliquiloculata | 1,223 |  | 159 | 1,064 |
| Pulleniatina praecursor | 1 |  |  | 1 |
| Pulleniatina primalis | 2 |  |  | 2 |
| Sphaeroidinella dehiscens | 546 |  | 37 | 509 |
| Tenuitella iota | 61 |  | 24 | 37 |
| Tenuitella parkerae | 1 |  |  | 1 |
| Trilobatus quadrilobatus | 7 |  | 7 |  |
| Trilobatus sacculifer | 1,622 | 1 | 355 | 1,266 |
| Trilobatus trilobus | 1,215 | 1 | 262 | 952 |
| Truncorotalia crassaformis | 932 | 3 | 256 | 673 |
| Truncorotalia crassula | 17 |  | 17 |  |
| Truncorotalia tosaensis | 4 |  | 1 | 3 |
| Truncorotalia truncatulinoides | 1,366 | 5 | 693 | 668 |
| Turborotalita cristata | 12 |  | 12 |  |
| Turborotalita humilis | 287 | 4 | 161 | 122 |
| Turborotalita quinqueloba | 1,277 | 276 | 667 | 334 |

Table S2: Number of assemblages sampled in the northern hemisphere (Northern) and the southern hemisphere (Southern), for each latitudinal zone, 0-30, 30-60, 60-90 degrees absolute latitude.

| **Latitudinal Zone** | **Northern** | **Southern** |
| --- | --- | --- |
| High | 372 | 1 |
| Mid | 600 | 324 |
| Low | 782 | 528 |

Table S3: The mean estimates for the change in thermal deviance if temperature increases by 1°C (red background) or decreases by 1°C (blue background), for each latitudinal zone, 0-30, 30-60, 60-90 degrees absolute latitude. The estimate is based on linear mixed effect models (‘mixed’) and first-order autoregressive models (‘autoregressive’). Values in square brackets indicate the 95% confidence interval. Beta values are positive when the magnitude of change in estimated sea surface temperature is greater than the magnitude of change in bio-indicated temperature.

| **Latitudinal Zone** | **Beta Coefficient_mixed_** | **Beta Coefficient_autoregressive_** |
| --- | --- | --- |
| High | 3.53 [2.12, 4.95] | 3.63 [0.24, 1.77] |
|  | 0.36 [0.06, 0.67] | 0.31 [0.03, 0.45] |
| Mid | 0.82 [0.05, 1.58] | 0.72 [-0.05, 1.57] |
|  | 0.09 [-0.24, 0.42] | 0.41 [-0.61, 0.52] |
| Low | 0.24 [-0.04, 0.53] | 0.14 [-0.16, 1] |
|  | 2.66 [1.56, 3.76] | 3.63 [1.77, 4.24] |

Table S4: The mean estimates for the assemblage turnover if temperature increases by 1°C (red background) or decreases by 1°C (blue background), for each latitudinal zone, 0-30, 30-60, 60-90 degrees absolute latitude. The estimate is based on chi-squared coefficients (‘chi-squared) and Bray-Curtis dissimilarity indices (‘bray-curtis’). Values in square brackets indicate the 95% confidence interval.

| **Latitudinal Zone** | **Beta Coefficient_chi-squared_** | **Beta Coefficient_bray-curtis_** |
| --- | --- | --- |
| High | -0.2 [-0.42, 0.02] | -0.02 [-0.1, 0.05] |
|  | -0.38 [-0.6, -0.16] | -0.11 [-0.18, -0.03] |
| Mid | 0.11 [0.05, 0.16] | 0.02 [0.01, 0.03] |
|  | -0.34 [-0.45, -0.24] | -0.02 [-0.03, 0] |
| Low | 0.13 [-0.19, 0.45] | 0.04 [-0.02, 0.1] |
|  | 0.05 [-0.17, 0.26] | 0.03 [0, 0.06] |

Table S5: The mean estimates for changes in species richness and extirpation probabilities if temperature increases by 1°C (red background) or decreases by 1°C (blue background), for each latitudinal zone, 0-30, 30-60, 60-90 degrees absolute latitude. Values in square brackets indicate the 95% confidence interval.

| **Latitudinal Zone** | **Richness** | **Extirpation** |
| --- | --- | --- |
| High | 0.23 [-1.05, 1.51] | -0.68 [-1.52, -0.03] |
|  | 0.63 [0.19, 1.06] | 0.07 [-0.32, 0.5] |
| Mid | 0.22 [-1.01, 1.45] | -0.18 [-0.45, 0.01] |
|  | 1.38 [0.29, 2.47] | 0.45 [-0.1, 1.31] |
| Low | -2.02 [-5.57, 1.54] | 0.11 [-0.39, 0.51] |
|  | 1.38 [0.29, 2.47] | -0.83 [-1.55, -0.09] |
